# Evolutionary Sparse Learning for phylogenomics

**DOI:** 10.1101/2021.07.19.452974

**Authors:** Sudhir Kumar, Sudip Sharma

## Abstract

We introduce a supervised machine learning approach with sparsity constraints for phylogenomics, referred to as evolutionary sparse learning (ESL). ESL builds models with genomic loci—such as genes, proteins, genomic segments, and positions—as parameters. Using the Least Absolute Shrinkage and Selection Operator (LASSO), ESL selects only the most important genomic loci to explain a given phylogenetic hypothesis or presence/absence of a trait. ESL does not directly model conventional parameters such as rates of substitutions between nucleotides, rate variation among positions, and phylogeny branch lengths. Instead, ESL directly employs the concordance of variation across sequences in an alignment with the evolutionary hypothesis of interest. ESL provides a natural way to combine different molecular and non-molecular data types and incorporate biological and functional annotations of genomic loci directly in model building. We propose positional, gene, function, and hypothesis sparsity scores, illustrate their use through an example and suggest several applications of ESL. The ESL framework has the potential to drive the development of a new class of computational methods that will complement traditional approaches in evolutionary genomics. ESL’s fast computational times and small memory footprint will also help democratize big data analytics and improve scientific rigor in phylogenomics.

## Introduction

Rapid acquisition and assembly of multigene and genomic datasets have fast-tracked the discovery of natural patterns and processes underlying the diversity of form and function. Central to these successes are statistical and computational methods for comparative analysis of molecular sequences, needed for applications ranging from building the tree of life to discovering the genomic loci underlying functional evolution. These methods have revolutionized data-driven discovery and hypothesis testing (BBSRC 2020; Kulathinal et al. 2020). The growth of data has resulted in a tsunami of information, putting data-driven biological research on steroids. With such increasingly bigger datasets, the pattern-matching paradigm of machine learning is poised to become a useful approach, complementing computational molecular evolution approaches based on stochastic models and evolutionary process descriptions (Nei and Kumar 2000; Yang 2014).

Here, we introduce supervised machine learning with a sparsity constraint for molecular evolutionary analysis (Wrinch and Jeffreys 1921; Tibshirani 1996; Ye and Liu 2012). We refer to this framework as evolutionary sparse learning (ESL). This approach treats the process of learning similarly to model selection with genetic loci, including genes, proteins, exons, introns, intergenic regions, individual genomic positions, and many other possibilities, as parameters. This feature contrasts with classical statistical methodologies in which sophisticated mathematical models describe substitutional and evolutionary processes to estimate parameters of these models, branch lengths, and sequence phylogeny to identify functionally important genomic loci and reconstruct evolutionary relationships of sequences.

By applying sparse learning directly to a multiple sequence alignment, ESL identifies the most important genetic loci (parameters) that predict a phylogenetic hypothesis or the presence or absence of a trait of interest. Alternative models involving different combinations of model parameters are automatically compared, and the model requiring the fewest loci (the “ sparse” solution) but offering the highest predictive ability is preferred. Thus, this approach emphasizes building a model with the fewest parameters while maximizing the model fit, similarly to traditional molecular evolutionary analyses investigating genomic features that drive a phylogenetic inference or underlie a trait. However, ESL analysis does not necessitate estimating standard evolutionary parameters, such as branch lengths, substitution rates between nucleotides, or site-wise evolutionary rate variability.

The ESL framework introduced here is different from recent machine learning applications in ecology and evolution to classify species (Suvorov et al. 2020; Zou et al. 2020), accelerate maximum likelihood phylogeny inference (Azouri et al. 2021), detect genomic regions under selection (Schrider and Kern 2016; Sugden et al. 2018), identify the best-fitting model of substitutions (Abadi et al. 2020), and detect autocorrelation of evolutionary rates in a phylogeny (Tao et al. 2019). These applications mainly focus on classification and need to use synthetic (computer-simulated) datasets for building predictive models. In ESL, no synthetic data are used as the biological hypotheses or the traits of interest are provided by the investigators to build models with the most informative genomic loci through machine learning. Of course, these ESL models can be used to make predictions as well (explained below).

In the following, we first present the general ESL framework, define several biologically relevant sparsity scores and prediction metrics, and outline several useful potential applications of ESL. We also introduce some examples to demonstrate ESL’s pattern recognition approach and illustrate its modest computational demands that speed up large-scale phylogenomics.

### A General Framework for Evolutionary Sparse Learning

ESL builds a logistic regression model that best maps the input multiple sequence alignment (MSA, X) to the probability of output response categories Y (phylogeny or trait, **Fig. 1a**). The logistic regression model is *f*(Y) = ***Xβ* (Fig. 1b)**. The input X consists of p positions (columns) and S sequences (row), Y is the categorical states of rows in X, and *f*(Y) is the logit link of class probabilities derived from Y. The ***β*** is the column vector of logistic regression coefficients that specifies the importance of positions (features) in predicting the outcome (Qiao et al. 2017). ESL is supervised machine learning because the outcome is provided in Y during ESL modeling (**Fig. 1**).

**Figure 1.**
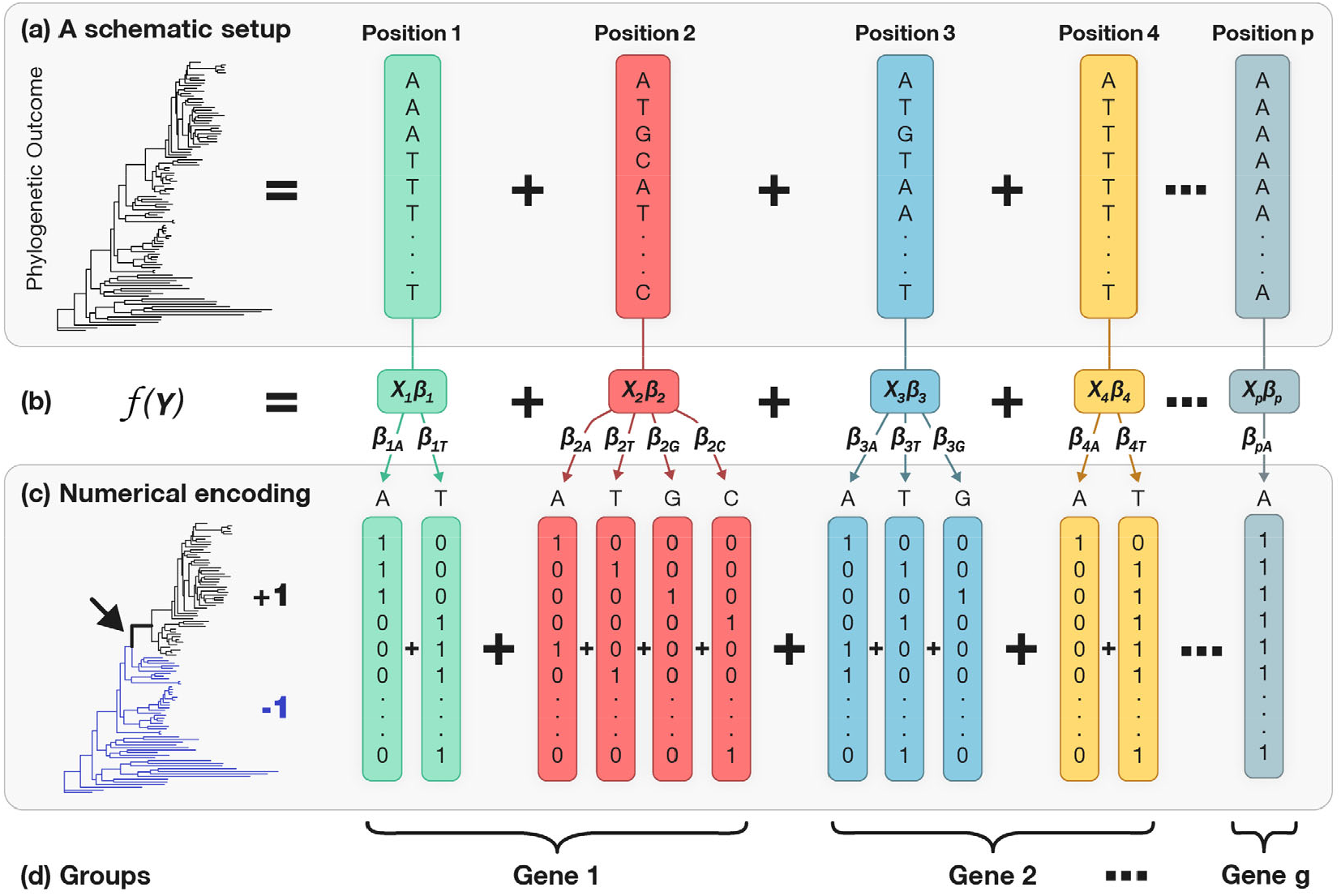
A schematic representation of models in Evolutionary Sparse Learning (ESL). (**a**) There are *p* positions in the sequence alignment, so the regression model (panel **b**) can contain as many as *p* variables, i.e., features in machine learning. The regression coefficient *β*_i_ is the degree of association between the base configuration at position *i* with the function of the outcome **Y**. The outcome is assigned to each sequence based on the phylogenetic relationship or the presence/absence of a trait. (**c**) One-hot encoding of the sequence alignment in which each base is represented by a column of bits. ESL estimates regression coefficient (*β*_*iw*_) for every bit-column (*w*) for every position *i*. In the response vector, all sequences belonging to the target clade (black) are represented by +1, and those in the other clade (blue) are represented by -1. (**d**) Positions can be clustered into groups (e.g., genes) for bi-level sparsity.

The learning part of ESL involves the estimation of importance (*β*_i_‘s). Phylogenomic datasets contain thousands of times more sites (positions) than the number of sequences (p ≫ S), but only a subset of these positions have substitutions that relate to the hypothesis. Therefore, a sparse solution of biological parameters (positions, genes, and loci) is usually appropriate to explain the phylogenomic hypothesis of interest. This process is related to feature selection in machine learning, which finds the optimal number of parameters (positions) for the model that minimizes the logistic loss.

A *l*_1_-regularized regression (Tibshirani 1996) (Least Absolute Shrinkage and Selection Operator, LASSO) accomplishes this task by minimizing the sum of the difference between the observed and the predicted output (e.g., logistic loss, *l*(*β*) in eq. 1) and the cost of including positions in the ESL model (overfitting penalty, second term in equation 1) (Tibshirani 2013).

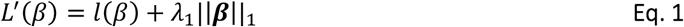

Here, the strength of association of the genetic variation at position *i* with the phylogeny is captured in the magnitude of the regression coefficients (*β*_i_’s), and ***β*** is a vector that contains all *β*_i_’s. Here, ∥***β***∥_1_ is defined as 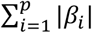.

We use logistic regression with ***l***_**1**_ regularization (Logistic lasso regression) because the outcome *Y* is a categorical variable. ***λ***_**1**_ is the regularization parameter that controls the sparsity of the model. This hyperparameter needs to be selected judiciously, as the choice of the regularization parameter controls the number of positions (loci) included in the model. When ***λ***_**1**_= 0, LASSO reduces to the standard statistical regression analysis that is known to suffer from the over-fitting problem (Meier et al. 2008). The choice of a large value of ***λ***_**1**_ will select a highly sparse model in which only a few (most important) positions will be included. That is, only a few positions will receive non-zero regression coefficient (*β*_i_ -=I 0) and, thus, a sparse solution will be generated (Hastie et al. 2015).

### Bi-level Regularization

Phylogenomic datasets are often short-and-fat, i.e., the number of sequences is hundreds to thousands of times smaller than the number of positions. This results in the curse of dimensionality^29^ because the number of model parameters (loci) is orders of magnitude larger than the number of sequences. Additionally, the logistic regression with mono-level sparsity does not produce a unique ESL model (unique solution) when *X* contains columns of a categorical variable (bit columns for genomic data, see below) (Meier et al. 2008; Tibshirani 2013; Hastie et al. 2015). The problem is alleviated somewhat by using LASSO with bi-level regularization. In bi-level regularization, columns (positions) are grouped into predefined groups, and the sparsity constraints are applied to groups and positions within groups. The bi-level sparsity is more practical in phylogenomics because we have biological annotations to cluster positions into groups. For example, individual genomic positions belong to genomic and functional groups, such as genes, exons, introns, intergenic regions, and other types of genomic segments. Furthermore, even non-contiguous positions may belong to the same group, e.g., groups of first codon positions in a codon alignment and other genomic annotations. Such information on groups of positions is used to impose a bi-level sparsity (Breheny and Huang 2009; Simon et al. 2013; Qiao et al. 2017). This is achieved by adding a penalty term that penalizes the inclusion of groups along with positions to accomplish a doubly sparse solution.

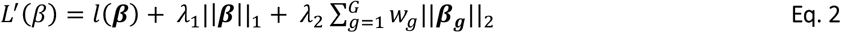

Here, *λ*_2_ is the group regularization parameter, ***β***_***g***_ is the vector of *β*_*i*_’s of positions belonging to group *g*, and ***w***_g_ is the weight assigned to group *g*. The norm ∥*β*_*g*_∥_2_ is defined as 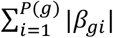, where *p*(*g*) is the number of positions in group *g* and *β*_*gi*_ is the regression coefficients for a position within group *g*. The second term in Equation 2 controls the sparsity for positions within the group. The third term controls the sparsity for groups. In the ESL framework, we use 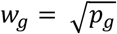 where *p*_*g*_ is the length of the group *g*, a common practice in sparse group lasso regression analysis (Zou and Hastie 2005; Simon et al. 2013; Hastie et al. 2015). Here a large value for *λ*_2_ will cause only a few groups to be included in the final ESL model and a large *λ*_1_ will allow only a few positions in each selected group to be retained. In phylogenomic datasets, we found small values of *λ*_1_ and *λ*_2_ to work well (see Fig. 2), but their selection needs to be done individually for each dataset (see later).

**Figure 2.**
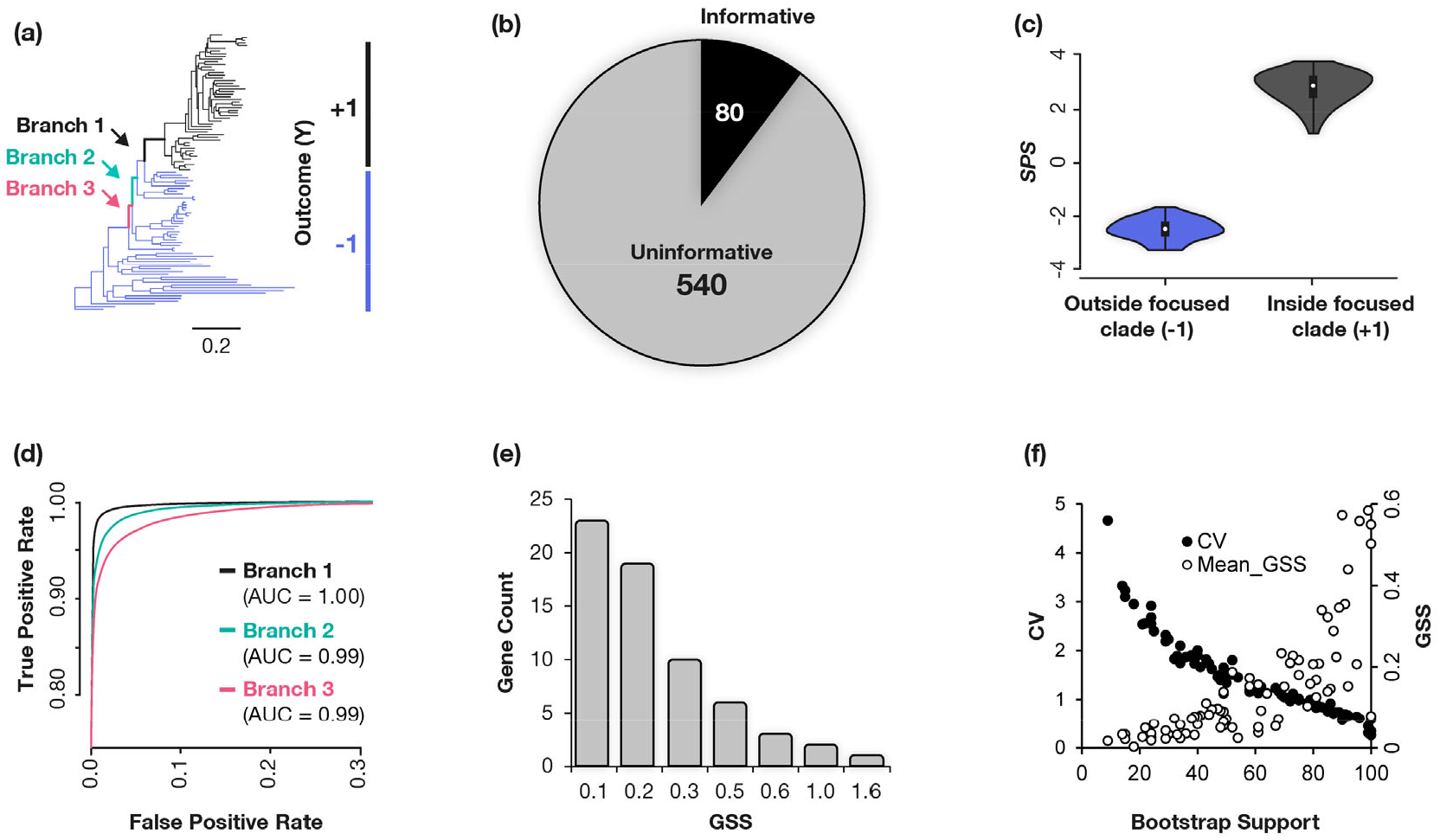
ESL analysis of a multiple sequence alignments of Plants species. **(a**) The Plant phylogeny with the sequences in one focused clade marked as +1 (orange) and the rest marked as -1 (blue). (**b**) Genes included in the ESL model after using sparse group lasso and the ridge regression for branch #1. (**c**) A violin plot showing the distribution of *SSS* for all the sequences using the ESL model for branch #1. (**d**) ROC curve showing the trade-off between the true positive rate and the false positive rate of classification of the ESL model for the phylogenetic partition induced by branch #1 (black). ROC curves of two other branches are also shown (blue and pink lines). ESL analyses were performed independently with similar settings for all three branches and ROC curves for different branches were calculated using genes that were selected in each ESL model. The areas under ROC curves (AUC) are also presented. (**e**) The distributions of *GSS* scores of genes included in the ESL model for branch #1. (**f**) Scatter plots showing the relationship between the proportion of bootstrap ESL models in which a gene appeared and its GSS (brown) and the coefficient of variation (CV) of *GSS*. ESL models were generated in the SLEP(Liu et al. 2011) software in MATLAB by analyzing a multiple sequence alignment of 620 genes (290,718 sites) from 103 Plant species. The “ sgLogisticR” function for bi-level logistic sparse group lasso regression was applied with starting feature regularization parameter (*λ*_1_ = 0.1) and group regularization parameter (*λ*_2_= 0.2). The square root of gene length was used as group weight. SLEP uses Moreau-Yosida Regularization algorithm and we specified 100 iterations to obtain an optimal parameter. In these iterations, the regularization parameters (*λ*_1_, *λ*_2_) were automatically optimized (0.0226 and 0.0129, respectively). We conducted Ridge regression analysis with *l*_2_ regularization, the “ gLogisticR” functions for genes selected in panel **b**. The regularization parameter *λ* = 0.1 was used and the squre root of gene length was used as group weight.

Statistically, bi-level sparsity is desirable because solutions are invariant under group-wise orthogonal re-parameterizations and statistically consistent when the number of groups to be discovered is small, i.e., a sparse solution is expected (Meier et al. 2008; Simon et al. 2013). The sparse solution is biologically realistic because only a (small) subset of positions and groups contain information for uniting a group of sequences when we wish to discover loci that are associated the most with a given biological outcome.

### Considering functional and biological categories

ESL analysis enables direct consideration of functional features of genes, such as GO annotations (Carbon et al. 2021). In this case, penalty terms are added in equation 2 to use a multi-level lasso, e.g., (Lozano and Świrszcz 2012; Qiao et al. 2017). In fact, genomic loci may belong to categories with overlapping compositions (e.g., the same gene belonging to multiple functional categories), and categories may even have hierarchical relationships. Such considerations would enable the direct discovery of important functional categories in ESL models, different from *post hoc* gene enrichment approaches frequently employed in evolutionary and functional genomics. In addition, a tree-structured group lasso is also feasible in ESL to relax the common assumption of independence among loci and groups (Liu and Ye 2010; Qiao et al. 2017).

### Numerical representation of the input data and response

The process of model selection requires numerical representations of input data and evolutionary outcomes. For input MSA (X), one-hot encoding is a common practice in machine learning. In this approach, as many binary columns represent each aligned position as the number of different bases found at that position across sequences in MSA (**Fig. 1c**). The one-hot encoding in machine learning is reminiscent of bit-wise representations used in molecular evolutionary analysis software (e.g., MEGA (Kumar et al. 1993)) for efficient implementation of Fitch’s parsimony algorithm (Fitch 1971) that requires logical operations such as intersections [ANDs] and unions [ORs]) to generate the most parsimonious count of substitutions required at a position given a phylogeny.

The maximum number of bits (bit-columns) required by any MSA position in **X** is four for nucleotides and 20 for amino acids. One may encode alignment gaps as their own bit-column if the presence/absence of gaps is meaningful. In addition, multi-base nucleotides and amino acid states in the alignments may be encoded in bit-columns of their constituent bases (e.g., nucleotides A and G bits for R). Ultimately, one-hot encoding transforms a sequence alignment **X** into a computer-friendly numerical format containing **c** bit-columns and **S** rows. Each column maps to exactly one position in this matrix, and multiple columns may represent a position. Generally, we reduce the memory requirements by preprocessing the **S x c** matrix, particularly by excluding all monomorphic bit-columns.

In ESL analysis, one-hot encoded data from different data types can be directly combined for the same set of organisms. For example, one-hot encoded amino acid and nucleotide MSAs can be used together by simple concatenation to build a super-matrix **X**. By one-hot encoding data such as the presence or absence of genes or other types of genomic annotations—such as methylation, post-translational modifications, and breakpoints—we can naturally combine heterogeneous datasets. Also, other molecular and non-molecular characteristics of organisms can be one-hot encoded and added to the input data matrix. Each of these disparate data types can be assigned its own groups, enabling ESL’s model-building to automatically compare the relative importance of different data types and even subgroups within each data type (e.g., nucleotide versus amino acid positions).

### Numerical representation of the response

In ESL, each sequence ***j*** in **X** needs to be associated with an outcome state (*y*_**j**_). **Y** is a column vector containing S binary outcome elements (**Fig. 1c**). We can set ***y***_**j**_ **= +1** for sequences with a given attribute (e.g., belonging to a cluster) and ***y***_**j**_ **= −1** for those without that attribute. Numerical values represent different categories, and their absolute values are chosen for computational convenience and interpretation. Beyond binary outcomes, computational approaches exist for sparse learning with multi-state outcomes (multi-class LASSO) and simultaneous consideration of multiple outcomes (multi-label LASSO); for a review, see (Liu et al. 2011; Chen et al. 2019). In a later section, we present different ways to specify the responses (Y) to conduct a range of evolutionary analyses.

### Regularization, stability selection, and class balance

In ESL, proper *regularization* is needed to obtain stable results. Therefore, one may use stability selection through a subsampling approach to make results robust to the regularization parameters chosen (Meinshausen and Bühlmann 2010). Stability selection yields finite sample family-wise error control and makes results robust to selecting regularization parameters. Cross-validation is another widely used approach for selecting optimal regularization parameters (Roberts and Nowak 2014). In cross-validation, the dataset is split into *k* independent subsets of sequences. The regression models are fitted to k -1 subsets and a wide range of regularization parameter values. The “ left-out” subset is used to validate the choice of regularization parameter values based on prediction error and repeat these steps multiple times. We selected the regularization parameter value for which the model has the lowest average prediction error.

Moreover, machine learning is most effective with balanced datasets, such that the number of sequences/species with and without the given trait is the same. In ESL, we use class weights, up-sampling with replacement of the minority class, or down-sampling the majority class to achieve class balance (Lunardon et al. 2014; Fabish et al. 2019). In the example discussed below, we used weights based on the class size for class balancing (Liu et al. 2011). This approach has the same effect as the upsampling or down-sampling of sequences with a replacement.

### Robust estimation of regression coefficients

After using lasso approaches in equations 1 and 2 to build an ESL model, the Ridge regression (*l*_2_-norm) should be applied for a more reliable estimation of β’s for the selected parameters (Le Cessie and Van Houwelingen 1992; Vágó and Kemény 2006). We can also use lasso (*l*_1_-norm) and Ridge (*l*_2_-norm) penalties together during the model selection. One may use ElasticNet (Zou and Hastie 2005) to improve the assignment of similar **β** values for strongly correlated parameters (Demir-Kavuk et al. 2011) for greater biological realism and model selection.

For estimating standard errors and confidence intervals of β’s and their linear combinations (sparsity scores below), both parametric and non-parametric approaches can be used. The parametric tests have been developed based on the distributional assumption of a test statistic, e.g., covariance test statistic computed from |*β*| (Halawa and El Bassiouni 2000; Kyung et al. 2010). Non-parametric statistical methods are also available to test the significance of regression coefficients (Cule et al. 2011; Lockhart et al. 2014). For example, we suggest using a bootstrap approach in which sequences are resampled with replacement within each class to build 100 (or more) bootstrap replicate datasets. Then, the bootstrap support for a position or group can be calculated as the proportion of bootstrap ESL models in which that position or group appears. One could also build a bootstrap consensus model from all the replicates. From this bootstrap procedure, we can also calculate variances of the sparsity scores defined below.

### ESL scores for use in phylogenomics

We define sparsity and prediction scores for individual positions and groups and overall hypotheses, which are linear functions of ***β*_i_′*s***. These new scores are expected to be useful for biological discoveries in molecular phylogenetics and evolution.

*Bit Sparsity Score* (***BSS***) is the absolute value of *β* for the bit-column corresponding to a specific character state at a particular position in the MSA. The score also represents the strength of association between the particular base (or character state) with the outcome (e.g., hypothesis). A vast majority of bit-columns receive **BSS** = 0 in the ESL analysis because only a small fraction of sites are likely to experience substitutions related to a hypothesis.

*Position Sparsity Score* (***PSS***) is the sum of absolute values of *β*’s (BSS values) for all the bit-columns that map to that position in the ESL analysis. Positions with nucleotide or amino acid-base configuration across sequences with limited or no concordance with the given hypothesis (***Y***) receive a PSS = 0. A large PSS indicates a high correlation with the hypothesis of interest.

*Group Sparsity Score* (*GSS*) is the sum of PSS of all the positions belonging to that group: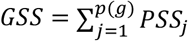, where *p*(*g*) is the number of positions in group ***g*** and *PSS*_*j*_ is the positional sparsity score for position *j*. Groups with limited or no ability to explain the specified hypothesis (outcome ***Y***) receive *GSS* = 0. A higher *GSS* indicates a group of positions that shows a strong relationship with the specified hypothesis.

*Functional Sparsity Score* (*FSS*) is the sum of sparsity scores for all *G* groups belonging to the given biological category: 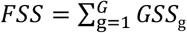. If a genomic locus (say **g**) belongs to multiple functional categories, it contributes to all relevant *FSS* scores. Importantly, all the FSS values are estimated from the same ESL analysis, so they have a common set of regularization parameters.

*Hypothesis Sparsity Score* (*HSS*) is the sum of sparsity scores for all *G* groups for the given hypothesis: 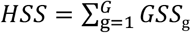. We expect it to be useful as an optimality criterion to discriminate among alternative phylogenetic hypotheses.

*Sequence Prediction Score* (*SPS*) is the predicted outcome (*Ŷ*) for a given sequence, which is computed by using the logistic lasso regression model.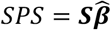, where 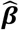 is a column matrix containing *β*’s, including the intercept of the regression model *β*_0_. ***S*** is the one-hot encoded vector for the sequence. Sequences with *SPS* greater than 0 are likely to belong to the class encoded by +1. In contrast, a value less than 0 makes the sequence likely to belong to the class coded by -1.

*Sequence Prediction Probability* (*SPP*) is the prediction probability that a sequence belongs to a class encoded by +1. The probability is computed as *SPP* = 1/(1 + *e*^-(*SPS*)^) for the logistic regression (Hosmer et al. 2013; Rao et al. 2016). One may use a probability ≥ 0.5 to classify a sequence into the class encoded by +1; otherwise, the sequence is classified in the class encoded by -1. To determine the best probability threshold for the highest accuracy level desired, we usually draw a receiver operating characteristic curve (ROC) (Fawcett 2006). ROC provides the relationship between the true-positive rate against the false-negative rate at different probability cut-offs (e.g., **Fig. 2d**). This can be done for the data used to build the model (training ROC) or during the cross-validation procedure.

### An Example Illustrating the Use of the ESL Framework

The ESL framework can be used to develop approaches for several types of applications. For example, we can determine positions and genes, along with their relative importance, in uniting a class of sequences (clade) in a given phylogenetic tree. In this case, *y*_*i*_ = +1 for sequences that belong to the given clade and -1 for the rest of the sequences in the MSA. Figure 2a shows a phylogeny of 103 Plant species (Shen et al. 2017) in which the clade of interest is assigned *y*_*i*_ = +1, and the sequences not in that clade are assigned *y*_*i*_ = -1. It corresponds to the branch #1, drawn with black lines, that partitions the tree into black and blue classes. The multiple sequence alignment consists of 290,718 sites that belong to 620 genes (Shen et al. 2017).

We applied the ESL framework to build a model that identifies the most influential genes that are likely to contain diagnostic substitutions. The ESL model was generated in the SLEP software in MATLAB (Liu et al. 2011). The “ sgLogisticR” function with bi-level logistic sparse group lasso regression was applied with starting feature regularization parameter (*λ*_1_ = 0.1) and group regularization parameter (*λ*_2_ = 0.2) that were selected by trial and error. These parameters are optimized iteratively during the model fitting step and coefficients (β’s) estimated (see **Fig. 2** legend). The square root of gene length was used as group weight. The ESL model contained only 80 genes, as *GSS* = 0 for the other 540 genes. Therefore, only 12.9% of genes were selected, which is a sparse solution (**Fig. 2b**). This sparsity is evolutionarily reasonable because not all genes are expected to have experienced a significant number of substitutions on any given branch in the phylogeny.

For these 80 genes, we used Ridge regression with *l*_2_ regularization to generate more accurate *GSS*, because *l*_1_ regularization is useful for initial model building (gene selection) and the *l*_2_ regularization yields more stable estimates of sparsity scores. This ESL model was used to estimate sequence prediction scores (***SPS***), which is useful to evaluate how well the ESL model can classify all the sequences used in building the model. We found that all the sequences in the black group received a positive *SPS*, whereas those in the blue group received a negative *SPS* in training (**Fig. 2c**). Therefore, the ESL modeling works well. The training ROC curve in **Fig. 2d** shows the trade-off between the true-positive (TP) and false-positive (FP) rates of sequence classification at different ***SPS*** thresholds. Based on the ROC curve, a TP rate of 100% and an FP rate of 0% are achieved with *SPS* = 0 (black curve). ESL models for phylogenetic partitions induced by other branches (#2 and #3; **Fig. 2a**) also showed very high classification accuracies, with the Area Under the Curve (AUC) greater than 0.99 (**Fig. 2d**). Overall, we found high training AUC for deep and shallow clades as well as for small and large clades (AUC > 0.99). Therefore, ESL prediction models are likely to be useful in placing a new sequence in the phylogenetic tree, an application that we are currently investigating.

The distribution of *GSS* values in the ESL model shows that a vast majority of genes receive a rather small sparsity score (**Fig. 2e**). Therefore, we used a bootstrap site resampling analysis (100 replicates) to identify genes that significantly contributed to the ESL model. Only seven genes appeared in more than 95% of the bootstrap ESL models. Thus, the final bootstrap-supported ESL solution was even more sparse and included only 1.2% of the genes. Similarly, only 0.6% of the positions (1,948) were included in more than 95% of the ESL models. The bootstrap support for a gene’s inclusion in the ESL model was highly correlated with its average *GSS* as well as the coefficient of variation (CV), which is the standard deviation of bootstrap estimates of *GSS* divided by the average bootstrap *GSS* (**Fig. 2f**). We also examined the frequency with which genes were included in ESL models built using datasets in which responses in Y were assigned +1 or -1 randomly. All genes were included in 10% - 68% of these ESL models, with a median of 39%. Therefore, random permutations of Y did not produce a statistically supported ESL model, as no genes were selected at a 95% significance level.

One well-known benefit of machine learning methods is their computational efficiency (time and memory) for high-dimensional datasets. This expectation is realized in the ESL analysis of the Plant dataset, which required *less than a minute* for all the analyses related to one branch on a personal desktop computer, consuming only 400 MB of computer memory for building any ESL model. The high computational efficiency of ESL meant that we could quickly apply ESL to all the nodes in this phylogeny and identify important node-specific loci. Further, ESL does not require the specification of within-group phylogenies when analyzing a specific branch (e.g., within blue and black clades, **Fig. 2a**). This is an interesting property because uncertainty regarding within-group phylogeny can create a need to integrate results over many alternative hypotheses in standard phylogenetic analysis, making them computationally very expensive for large datasets.

### Other applications of ESL analysis

In the above example, we partitioned all sequences in a phylogeny into two classes based on our interest in identifying genes and positions that are diagnostic of a group’s monophyly. In the interest of discovering positions that discriminate between sister clades, we can set ***y***_***i***_ = +1 for sequences in one clade (class) and -1 for the sequences in the other clade. In this case, we simply remove all other sequences from the MSA while building the ESL model. The outcome is a list of significant positions and corresponding genes in phylogenomic analyses in which the focus is on phylogenetic inference.

On the other hand, we seek a model in which positions or domains that distinguish paralogous genes are revealed if the divergence of two clades of interest corresponds to a gene duplication event in a multigene family sequence alignment. This would be useful in functional genomics investigations. In fact, the two classes of sequences in ESL analyses can be specified for any combination of clades, paralogs, and even sequences. So, it is very flexible. For example, we are currently investigating the effectiveness of ESL analysis to identify genes and functional categories in which gene evolution underlies the emergence of convergent traits.

We also envision a novel application of the ESL framework in which models are built under two contrasting evolutionary hypotheses. In this case, the relative importance of each locus that distinguishes these hypotheses can be determined based on the difference in their *GSS* for two hypotheses. We are exploring this idea to develop an approach to identify genes that may mislead phylogenomic inferences, which is quite common, e.g., (Shen et al. 2017; Walker et al. 2018).

### Similarities and Differences Between the ESL and Maximum Likelihood analyses

Statistically, the model building in ESL using equations 1 and 2 is equivalent to maximizing the product of the likelihood and a prior on the penalty (Breiman 1996; Tibshirani 1996; Fan and Li 2001; Breheny and Huang 2009). The regression coefficient, *β*, in ESL is comparable to the maximum *a posteriori* estimate when considering a Gaussian likelihood function and a Laplacian prior on *β*’s (Figueiredo 2002). In this case, 1*β*_m_1 *≥* |*β*_k_| means that bit-column *j* is as much or more correlated with the partial residual than position *k* (Frey 2018). The same applies to sparsity scores of positions that contain these two bit-columns, i.e., *PSS*_v_ *≥ PSS*_w_ where bit-column *j* is for position *v* and bit-column *k* is for position *w*. In the Maximum Likelihood analysis, |*ln*L_v_| *≥* |*ln*L_w_| means that the likelihood of position *v* is higher than position *w* for the given phylogenetic hypothesis (Felsenstein 1992). All sparsity scores defined here (*PSS, GSS, FSS*, and *HSS*) are linear sums of bit-wise sparsity scores (*BSS*), so they have analogous statistical interpretations.

However, there are many notable differences between ESL and ML. For example, ESL analysis does not use traditional substitution models that incorporate unequal rates of base substitutions, compositional bias, and heterogeneity of evolutionary rates and substitution patterns across positions. Nonetheless, we would be incorrect in stating that ESL analyses are agnostic to such biological features. For example, ESL does not assume that all positions in the alignment and all bases at a position follow the same evolutionary rate or have equal importance. Instead, the best ESL model assigns different weights to bases, positions, and groups, with many bases, positions, and groups receiving a zero weight for the given hypothesis. This is enabled by one-hot encoding that transforms MSA in which alternative bases at each position are separated into their binary columns.

Because of one-hot encoding, a composite of two-state models (one for each bit-column) describes a position rather than a single four-or 20-state substitution model in traditional analysis. This means that the same substitution model is not assumed for all the positions in the alignment or all the positions in a gene, unlike traditional methods. Moreover, the complexity of the model is a function of different base types found at each position. That is, position-by-position consideration of substitution matrices is intrinsic to ESL, but not in the same way or extent as in classical molecular phylogenetics. A major avenue of future research will be to investigate the relationship of ESL, theoretically and empirically, with Maximum Likelihood and other statistical methods in molecular evolutionary analysis. In particular, it will be interesting to test the robustness of ESL methods as compared to existing methods to the non-stationarity, non-reversibility, non-independence, and non-uniformity of substitution models across lineages and positions in multiple sequence alignments, as compared to traditional methods that tend to make these assumptions for analytical tractability.

## Conclusions

Overall, we expect ESL to complement existing methods of molecular evolutionary analyses because they serve different purposes. For example, one would need classical methods when the goal is to estimate branch lengths in a phylogeny, instantaneous rates of different types of mutations and substitutions, neutrality index, and the degree of heterogeneity of evolutionary rates among sites. They constitute fundamental properties of the evolutionary processes and natural selection, which are best estimated using statistical methods that model those properties. But, we may first use ESL to gain insights about evolutionary relationships and functional loci and then test them using currently standard statistical methods of computational molecular evolution. We also envision hybrid ESL approaches in which the input matrix contains estimates of such properties for genetic loci alongside the sequence alignments. Ultimately, we expect the utility of ESL to be limited only by one’s imagination, as it provides a flexible framework to construct approaches for *de novo* discovery and hypothesis-testing.

In conclusion, the power of machine learning in phylogenomics has only begun to be harnessed. ESL brings time-tested mature advances of sparse learning to phylogenomics. It provides a new way of conducting evolutionary analysis and enables a natural combination of heterogeneous datasets. Our simple example establishes the premise of ESL for developing methods for evolutionary analysis, which should motivate theoretical and computational investigations of the powers and pitfalls of ESL.

## Acknowledgments

We are grateful to Qiqing Tao for extensive technical assistance and comments on the manuscript. Prof. Slobodan Vucetic provided feedback on the theoretical properties of the sparse learning methods. Thanks are also due to Drs. Jack Craig, Jose Barba-Montoya, and Antonia Chroni for their many helpful suggestions to improve the manuscript. This research was supported by grants from the U.S. National Institutes of Health to S.K. (GM-0126567-01).

## Author Contributions

SK conceived ideas presented, conducted data analysis, and wrote the manuscript. SS implemented and advanced the ideas, performed the bulk of the data analysis, and co-wrote the manuscript.

## Data Availability and Code Availability

The dataset and source codes are available on GitHub. (https://github.com/ssharma2712/Evolutionary_Sparse_Learning_ESL).

